# An intron proximal to a PTC enhances NMD in *Saccharomyces cerevisiae*

**DOI:** 10.1101/149245

**Authors:** Jikai Wen, Muyang He, Marija Petric, Laetitia Marzi, Jianming Wang, Kim Piechocki, Tina McLeod, Anand K. Singh, Vibha Dwivedi, Saverio Brogna

## Abstract

Nonsense mediated mRNA decay (NMD) is regarded as the function of a specialized cytoplasmic translation-coupled mRNA decay pathway in eukaryotes, however, whether a premature translation termination codon (PTC) will lead to NMD often depends on splicing a downstream intron in the nucleus. Deposition of the exon junction complex (EJC) on mRNA is understood to mediate such splicing-dependent NMD in mammalian cells. The budding yeast, *Saccharomyces cerevisiae*, which has introns in only 5% of its genes, characteristically at the start of the coding region, and lacks proteins essential for EJC assembly, is not expected to undergo splicing-dependent NMD. However, we found that the presence of an intron near a PTC can also enhance NMD in this organism, regardless of whether it is downstream or upstream. These data provide evidence for a hitherto unsuspected EJC-independent mechanism linking translation and pre-mRNA in *S. cerevisiae*.

## Introduction

Nonsense-mediated mRNA decay (NMD) describes the observation that mutant or abnormally processed mRNAs that encode a premature translation termination codon (PTC) are typically at a low level and unstable in eukaryotes. NMD is primarily regarded as a quality control mechanism of gene expression that prevents wasteful production of potentially toxic truncated proteins by rapid destructing abnormal mRNAs. Additionally, by targeting functional mRNAs carrying specific features like an abnormally long 3’ UTR, NMD is understood to modulate protein expression in different cellular processes, including the cell stress response and some specific cell differentiation processes (Jaffrey and Wilkinson, 2018; Nasif et al., 2018). NMD was first branded as a specific process in *Saccharomyces cerevisiae* (Leeds et al., 1992); its mechanism and the roles that NMD play in gene expression have extensively been studied since, in this and in several other model organisms, as previously reviewed (Fatscher et al., 2015; He and Jacobson, 2015; Hug et al., 2016; Jaffrey and Wilkinson, 2018; Karousis and Muhlemann, 2019; Kurosaki et al., 2019; Lykke-Andersen and Jensen, 2015). NMD is coupled to translation and requires a set of evolutionarily conserved proteins, including UPF1, UPF2, and UPF3 (Lloyd, 2018). However, how the concerted action of NMD factors could form an mRNA surveillance complex capable of distinguishing PTCs from normal stop codons, either at or after translation termination, is unsatisfactorily understood in any organism. In the absence of a coherent mechanism and many conflicting observations it has been proposed that eukaryotes perhaps do not possess any such surveillance mechanism (Brogna et al., 2016). Moreover, recent evidence indicates that UPF1 plays broad roles in fundamental processes of gene expression, in the nucleus as well as in the cytoplasm, and these more than its requirement in NMD might explain UPF1 conservation in eukaryotes (Singh et al., 2019).

Specifically, current models cannot satisfactorily explain the link between pre-mRNA splicing and NMD observed in different organisms. Initial studies in human cells concluded that a PTC induces strong NMD more frequently when it is located upstream of an intron (Carter et al., 1996; Thermann et al., 1998; Zhang and Maquat, 1996; Zhang et al., 1998), suggesting that NMD primarily depends on pre-mRNA splicing in mammalian cells. Although later studies have downplayed the early view that NMD primarily depends on splicing in mammalian cells (Buhler et al., 2006; Eberle et al., 2008; Silva et al., 2008; Singh et al., 2008), a downstream intron remains a strong genome-wide predictor of whether a PTC will lead to NMD in mammalian cells (Hurt et al., 2013; Lindeboom et al., 2016; Mendell et al., 2004). Deposition of the exon junction complex (EJC) is understood to mark a splice junction in mammalian cells during splicing in the nucleus until the EJC is removed by a translating ribosome in the cytoplasm (Le Hir et al., 2000a; Le Hir et al., 2000b; Le Hir et al., 2016). Current models predict that in mammalian cells one or more downstream EJC distinguishes PTCs from normal stop codons and acts as the signal for NMD activation, whilst normal stop codons, since they are typically in the last exon have no downstream EJCs that could activate NMD (Maquat, 2004; Nagy and Maquat, 1998). It has also been reported that a downstream intron and EJC deposition might enhance NMD in evolutionarily distant organisms, including *Neurospora crassa* and plant species (Nyiko et al., 2013; Zhang and Sachs, 2015).

Notably though there are examples of splicing-dependent NMD that the EJC model cannot explain (Brogna and Wen, 2009; Buhler et al., 2006; Carter et al., 1996; Wang et al., 2002). Specifically, splicing can enhance NMD in *Schizosaccharomyces pombe* mutant strains depleted of core EJC proteins (Wen and Brogna, 2010), and EJC deposition is not likely to be the reason why normally-occurring mRNAs with one or more exon-exon junctions in the 3’UTR are affected by NMD in the protozoan *Tetrahymena* in mutant lacking EJC core components (Tian Miao, 2017). It is therefore conceivable that there is a mechanism other than EJC deposition that links splicing to NMD in these and other organisms.

Although the genome of *S. cerevisiae* does not code for Y14 and MAGO, in view that splicing-dependent NMD might occur independently of EJC deposition, we wanted to investigate whether splicing could also affect NMD in this much-studied organism in which both NMD and pre-mRNA splicing have been extensively characterised. Current models predict that the key NMD determinant in *S. cerevisiae* is an abnormally long distance between the PTC and the mRNA 3’ end, which possibly prevents an interaction of poly(A) binding protein (PABPC) with the terminating ribosome that is required for normal translation termination (He and Jacobson, 2015). Only a few protein-coding genes contain introns in *S. cerevisiae* (285 out of 5749, based on a recent study (Juneau et al., 2006)) and the vast majority of these (∼97%) have a single intron located close to the start of the coding region (Schreiber et al., 2015). However, intron-containing genes include those for ribosomal proteins and other highly expressed proteins, hence the rate of pre-mRNA splicing is globally high in this organism, accounting for over 24% of mature cellular mRNAs (Ares et al., 1999; Juneau et al., 2006; Kupfer et al., 2004; Lin et al., 1985). Inefficient splicing, alternative splice site selection, or splicing saturation caused by a high transcription rate, frequently produce unspliced transcripts; which, because they carry intron-encoded PTCs distant from the normal 3’UTR, can induce strong splicing-independent NMD (He et al., 1993; Kawashima et al., 2014; Kawashima et al., 2009; Sayani et al., 2008).

We examined the mRNA levels and stabilities of various gene-reporters with an intron either upstream or downstream of PTC. These data clearly indicate that splicing can enhance NMD in either context, as far as the intron is proximal to the PTC, and indicate the existence of an EJC-independent mechanism that links pre-mRNA splicing to translation in *S. cerevisiae*.

## Results

### Splicing enhances NMD in *S. cerevisiae*

We assessed whether splicing affects NMD in *S. cerevisiae* by examining the expression of GFP NMD reporters with or without an intron (Figure 1A, and Material and Methods). A PTC was introduced at codon position 6 (PTC6) or 140 (PTC140); PTC6, being early in the coding region, is expected to induce strong NMD, while PTC140, being in the second half of the GFP ORF (58% into the CDS), should be less affected based on earlier studies (Kuperwasser et al., 2004; Wen and Brogna, 2010). Three introns, derived from the *ACT1, CYH2* and *UBC4* genes, were examined; all carry canonical splice signals and are efficiently spliced both in vitro and in vivo (Abelson et al., 2010; Lesser and Guthrie, 1993; Swida et al., 1986). The intron-containing reporters, in which the intron was inserted at codon position 110 (Figure 1A), are designated by the suffixes A-ivs, C-ivs and U-ivs, carrying the *ACT1, CYH2* and *UBC4* introns, respectively.

**Figure 1.**
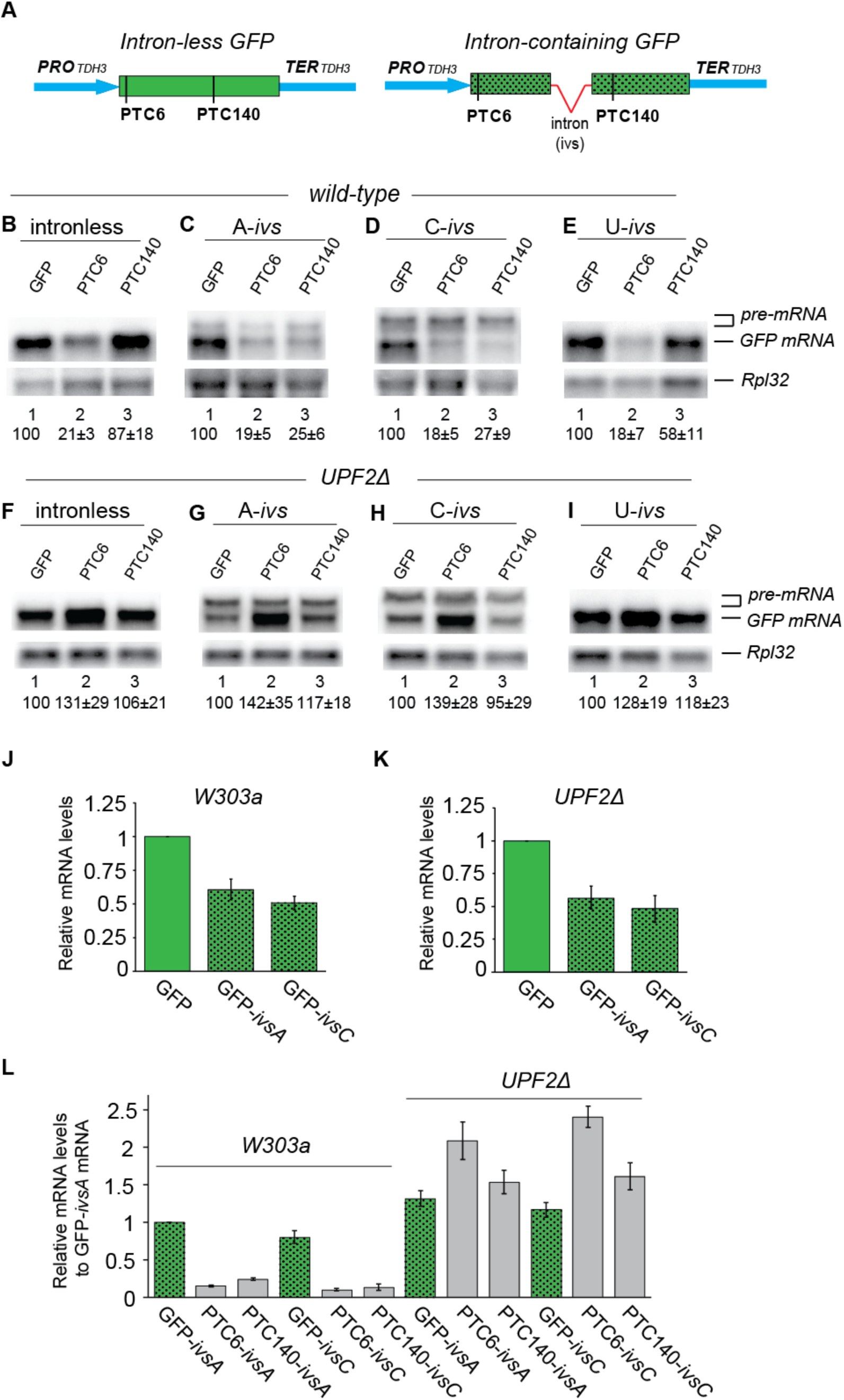
Splicing enhances NMD in *S. cerevisiae*. (**A**) Schematic maps of NMD reporters. The GFP gene was cloned into pRS304, which is regulated by the promoter and terminator of *S. cerevisiae TDH3* gene. The schematic of the intron-less constructs is on the left, while on the right is that of the intron-containing constructs carrying one of the three *S. cerevisiae* introns, labeled *ivs*; A-*ivs*, C-*ivs* and U-*ivs*, which derive from *ACT1, CYH2* and *UBC4* genes, respectively. (**B-E)** Northern blot analysis of mRNA levels in wild-type (W303a) cells transformed with the indicated NMD reporters. Top panels show hybridization with a GFP-specific probe, the bottom panel shows a probe specific for the ribosomal protein L32 mRNA (Rpl32), as a loading control. The values below each lane are percentages with standard deviations of the level of GFP mRNA relative to that of the corresponding PTC-less control, lane 1 in each panel. Band intensities were quantified using a phosphorimager and divided by the RpL32 band intensity of the same blot (see Materials and methods). (**F-I**) Similar Northern blot analysis carried out in *UPF2Δ.* Quantifications are based on three biological repeats. (**J-K**) RT-PCR quantification of GFP-ivsA and GFP-ivsC mRNA levels relative to that of the intron-less GFP reporter, in either wild-type *W303a* (J), or *UPF2Δ* (K). (**L**) Relative mRNA levels of different NMD reporters in wild-type and *UPF2Δ*. The mRNA levels were normalized by that of 18S rRNA (assessed in parallel using a 1/100 dilution of the same cDNA sample) and compared to that of GFP-ivsA in wild-type defined as 1. Details about RT-PCR quantification are in Supplementary Figure 1.

All PTC-less reporters produced intense GFP fluorescence comparable to the intron-less reporter (Supplementary Figure 1A), showing that all three introns are substantially correctly spliced. As expected, no fluorescence was detected in the strains transformed with the PTC-containing (PTC+) reporters (Supplementary Figure 1A and data not shown). Transcripts of the expected sizes were detected by Northern blotting: a single band corresponding to mature mRNA of the intron-less reporter and an additional weaker band corresponding to unspliced transcripts of two of the intron-containing reporters, (Figure 1B-1E). The unspliced transcripts were visible in the A-ivs and C-ivs reporters (we have termed these “pre-mRNAs” to distinguish them from the non-spliced mRNAs produced from the intron-less reporters that do not undergo splicing). Although the pre-mRNA accumulation indicates that these two introns are not always fully spliced, unlike U-ivs, the spliced mRNA bands were apparent and more intense in both cases, indicating that they are more often correctly spliced. Only the two expected fragments corresponding to either the spliced mRNA or pre-mRNA were also detected by RT-PCR (Supplementary Figure 1B), and no evidence of alternative splice site usage was observed by sequencing clones of a RT-PCR product spanning the entire coding region of the spliced mRNAs (data not shown). Real-time RT-PCR quantification, using a primer pair that amplifies only the spliced transcript since one of the primers spans to the splice junction (Supplementary Figure 1C-D), confirmed that A-*ivs* and C-*ivs* PTC-less reporters are well-expressed and produce 50-60% of mRNA compared to the intron-less version (Figure 1J and 1K).

Regarding NMD, the Northern blots show that PTC6 strongly reduces mRNA levels (∼20% of PTC-less control, regardless of the presence of an intron, as expected (Figure 1B-1E). In contrast, while PTC140 causes only a minor mRNA reduction of the intron-less reporter (87%), a strong reduction is apparent in the reporter carrying either A-ivs (25%) or C-ivs (27%) (Figure 1C and D), and a moderate reduction in the U-ivs reporter (58%) (Figure 1E). The data indicate that particularly the longer A-ivs and C-ivs intron enhance NMD. These conclusions can be drawn from pair-wise comparisons of mRNA levels between PTC and PTC-less versions of otherwise similar constructs, with or without an intron at a given position. As the variable in each of the pairs is the presence or absence of a PTC, it is unlikely that factors other than the PTC, such as changes in splicing efficiency, transcription, and potentially other processes, could have affected these mRNA levels. Furthermore, the levels of PTC+ mRNAs are higher in *UPF2Δ* (Figure 1F-1I and 1L) and *UPF1Δ* (Supplementary Figure 1F-1I), as expected, due to NMD suppression. Further real-time RT-PCR quantifications of steady-state mRNA levels of the different A-ivs and C-ivs reporters confirmed that the presence of a proximal intron enhances NMD in the wild-type strain and that mRNAs levels are increased in *UPF2Δ* (Figure 1L).

We also assessed decay kinetics of the different mRNAs following transcription inhibition in WT and *UPF2Δ* (Figure 2). The data confirm that both PTC140-ivsA and PTC140-ivsC transcripts are at lower levels at steady-state and decay faster than the intron-less version in wild-type, with kinetics similar to the PTC6-ivs mRNA that is strongly affected by NMD (Figure 2A-C). The PTC+ mRNA levels, and those of the intron-containing reporters in particular, mostly go down within the first 10 minutes from transcription inhibition, while they are more stable at later time points, suggesting that transcripts are perhaps most susceptible to NMD for a limited time after they have been synthesized, similar to what has been observed by some studies in mammalian cells, Drosophila and *S. pombe* (Gatfield et al., 2003; Trcek et al., 2013; Wen and Brogna, 2010). All transcripts show similar decay kinetics in *UPF2Δ*, as expected from NMD suppression. Notably, the steady-state levels of PTC+ mRNAs (most strikingly PTC6) were even higher than the corresponding PTC-less controls in the NMD mutants (Figure 1F-1I and 1L, Figure 2D-2F and and Supplementary Figure 1F-1I; additional examples are shown further below in Figure 3C-2E). This puzzling effect has previously been observed in *S. cerevisiae* and other organisms (Eberle et al., 2008; Gatfield et al., 2003; Metzstein and Krasnow, 2006; Peltz et al., 1994; Wen and Brogna, 2010); its significance has previously been discussed (Brogna et al., 2016; Brogna and Wen, 2009; Wen and Brogna, 2010).

**Figure 2.**
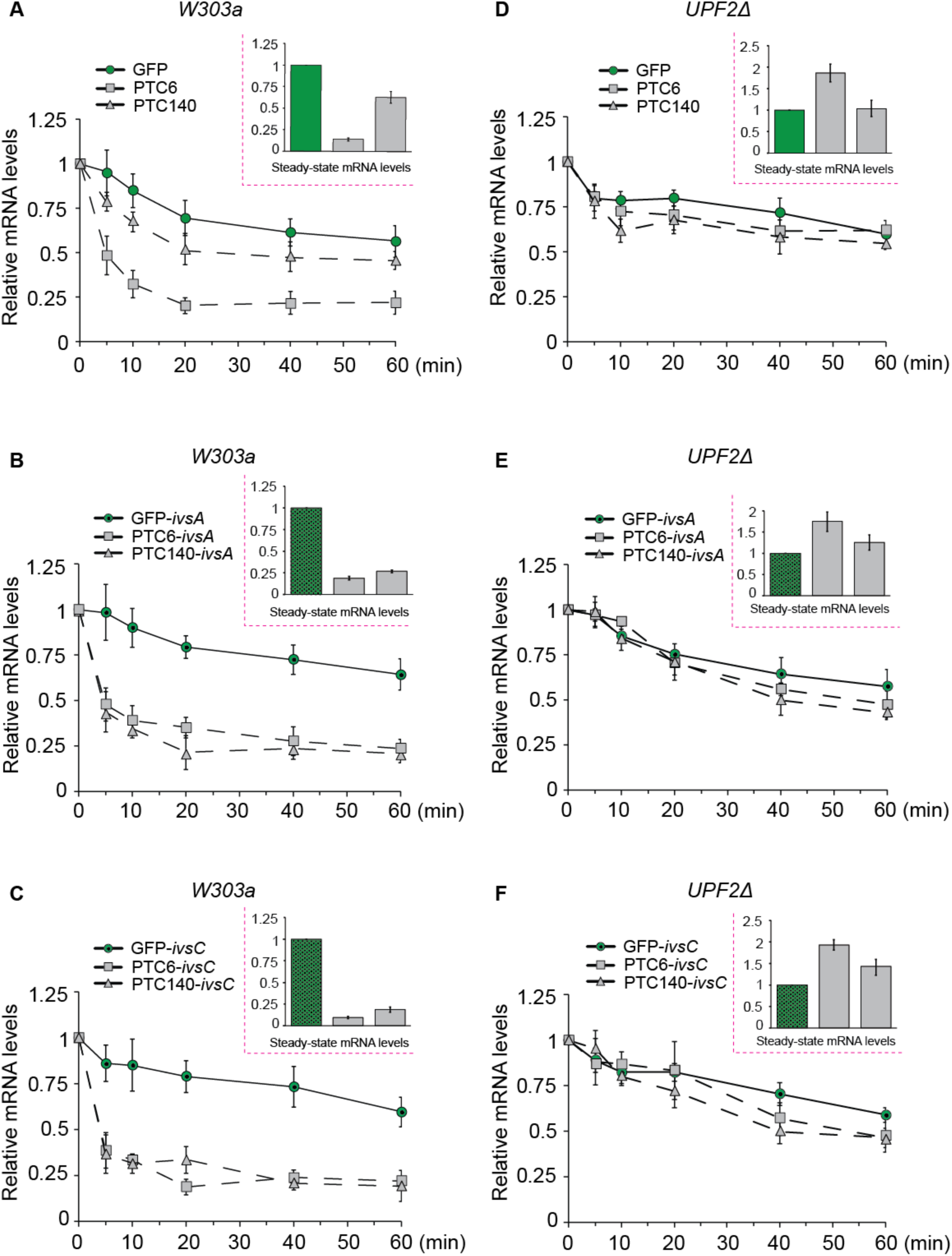
Splicing-dependent NMD correlates with a faster mRNA turnover. (**A**) Line profiles show mRNA levels of the intron-less reporters indicated at specific time points after transcription inhibition in wild-type (*W303a*) (0, 5, 10, 20, 40 to 60 min, see Material and Methods). Inset shows steady-state mRNA levels of the reporters. All mRNA levels were quantified by real-time RT-PCR using a primer pair specific for the spliced mRNA or intron-less construct (see Material and Methods and Supplementary Figure 1). (**B**) Same analysis as in A for the *ACT1* intron-containing reporters (*ivsA*). (**C**) Same as above for the *CYH2* intron-containing reporters (*ivsC)*. (**D-F**) Same analysis of the reporters described in A-C expressed in the *UPF2Δ* strain. All the measurements were repeated three times starting from independent cultures.

**Figure 3.**
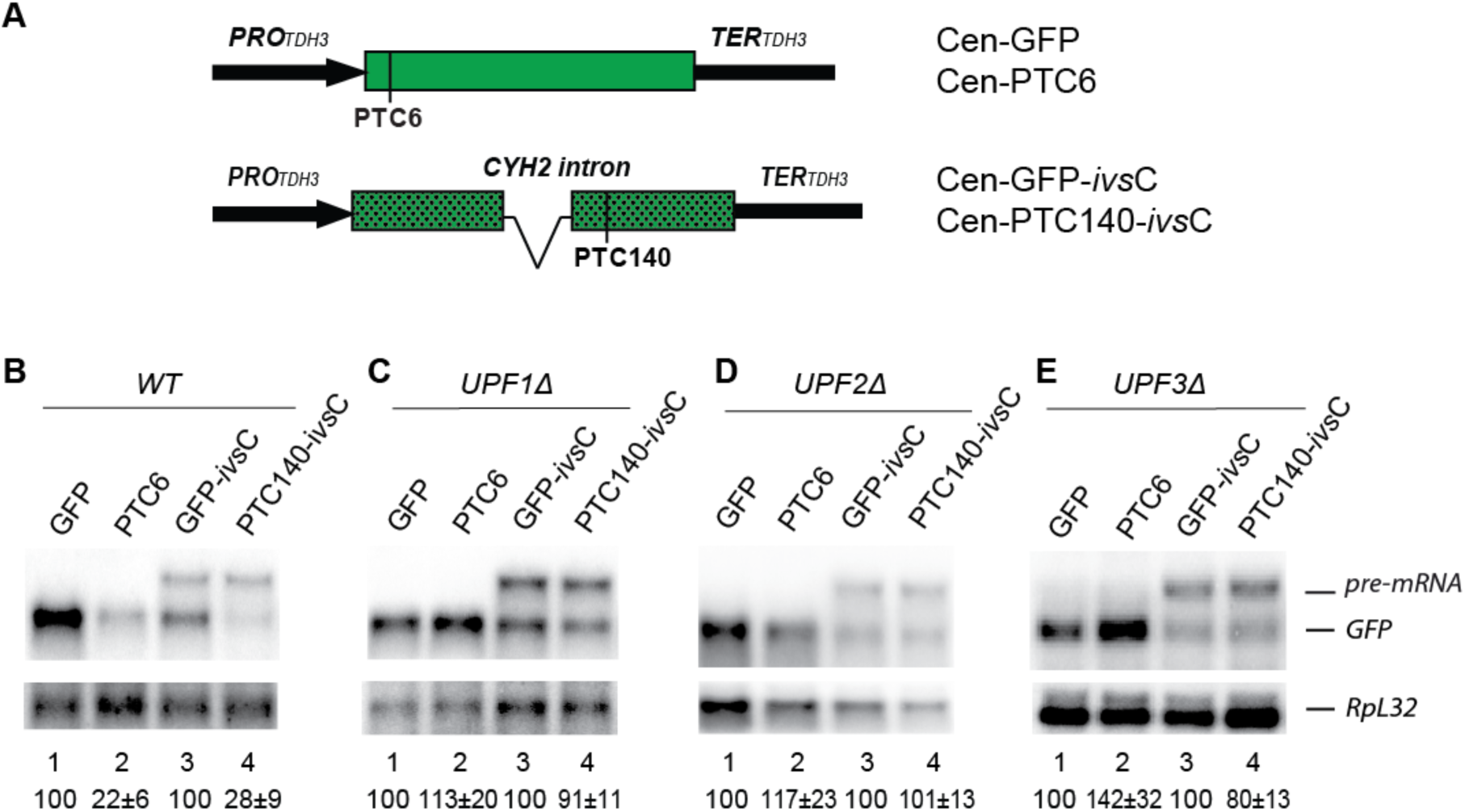
Splicing-dependent NMD is independent of reporter expression level. (**A**) Schematic maps of centromeric NMD reporters. (**B-E**) Northern blot analysis of reporter expression carried out in wild type (*BY4742)* (B), *UPF1Δ* (C), *UPF2Δ* (D), *UPF3Δ* (E). Top panels show the GFP hybridization signal; the bottom panels show those of the normalization control *Rpl32B* mRNA. Quantifications are based on three biological repeats.

The A-ivs or C-ivs reporter pre-mRNAs also appear to be affected by NMD as their levels increased in *UPF2Δ* (Figure 1G-H and Supplementary Figure 1F). Retention of either the A-ivs or C-ivs intron results in a frameshift that introduces a PTC either nine or five bases respectively, downstream of the 5’ ss. These PTCs are at a distance from the 3’UTR either similar (A-ivs) or further (∼200 nt greater in C-ivs) than that of PTC6 in the initial intron-less reporter. Hence, these PTCs were expected to induce strong NMD even in the absence of an upstream PTC.

Splicing-dependent NMD is independent of expression level, as similar observations were made using different strains (BY4742 wild-type strain, UPF2Δ, UPF2Δ and UPF2Δ from the Yeast MATalpha collection - Supplementary Table 1) transformed with centromeric plasmid constructs expressing either wild-type GFP or GFP containing a PTC, in the presence or absence of the C-ivs intron (Figure 3A) – vs. those described above that are 2μ-based plasmid constructs. The intron-less construct carrying PTC6 served as a splicing-independent NMD reporter, while PTC140-Civs served as a splicing-dependent NMD reporter. A pre-mRNA band is apparent with the latter, indicating that the intron is not always spliced in this position, as expected from the results with corresponding 2μ reporters, yet a band corresponding to the spliced transcript is clearly apparent, and its level is drastically reduced compared to the PTC-less counterparts in the wild-type (Figure 3B) but not in the three upf mutants, consistent with NMD suppression (Figure 3C-3E).

### Intron deletion inhibits NMD of endogenous intron-containing genes

To examine the effect of splicing on NMD further, we generated additional reporters derived from two endogenous intron-containing genes, *CYH2* (*RPL28*) and *RPL11B* (Figure 4). A PTC was introduced either 54 nt downstream (*CYH2*) or 30 nt upstream of the intron (*RPL11B*), and the corresponding intron-less constructs were generated via cDNA cloning (Figure 4A). To distinguish the reporters from the endogenous transcripts, as well as to inhibit splicing-independent NMD at these otherwise early PTCs, a 300 bp fragment was inserted in-frame at the start of the coding region of each gene. We found that the PTC induced apparent NMD in both genes. The levels of both PTC+ spliced mRNAs were reduced to ∼25% relative to the corresponding PTC-less controls, when assayed either by quantitative RT-PCR (Figure 4B and 4C), or Northern blotting (Figure 4D, lanes 1-2 and 5-6). However, no evidence of NMD was detected in the corresponding cDNA-derived reporters lacking the intron (constructs labeled with the *Δivs* suffix) neither by RT-PCR (Figure 4B and 4C) nor by Northern blotting (Figure 4D, lanes 3-4 and 7-8). It appears that these late PTC induce NMD only in the presence of an intron, similar to what we reported above for the GFP reporters, irrespective of whether the PTC is located downstream (as in *CYH2*) or upstream (as in *RPL11B*) of the intron. All PTC+ mRNA levels are increased, either comparable to the corresponding PTC-less controls in *UPF2Δ*, or as discussed above for previous reporters, even at higher levels (Figure 4B and 4C bottom panels, and Figure 4E), consistent with NMD suppression.

**Figure 4.**
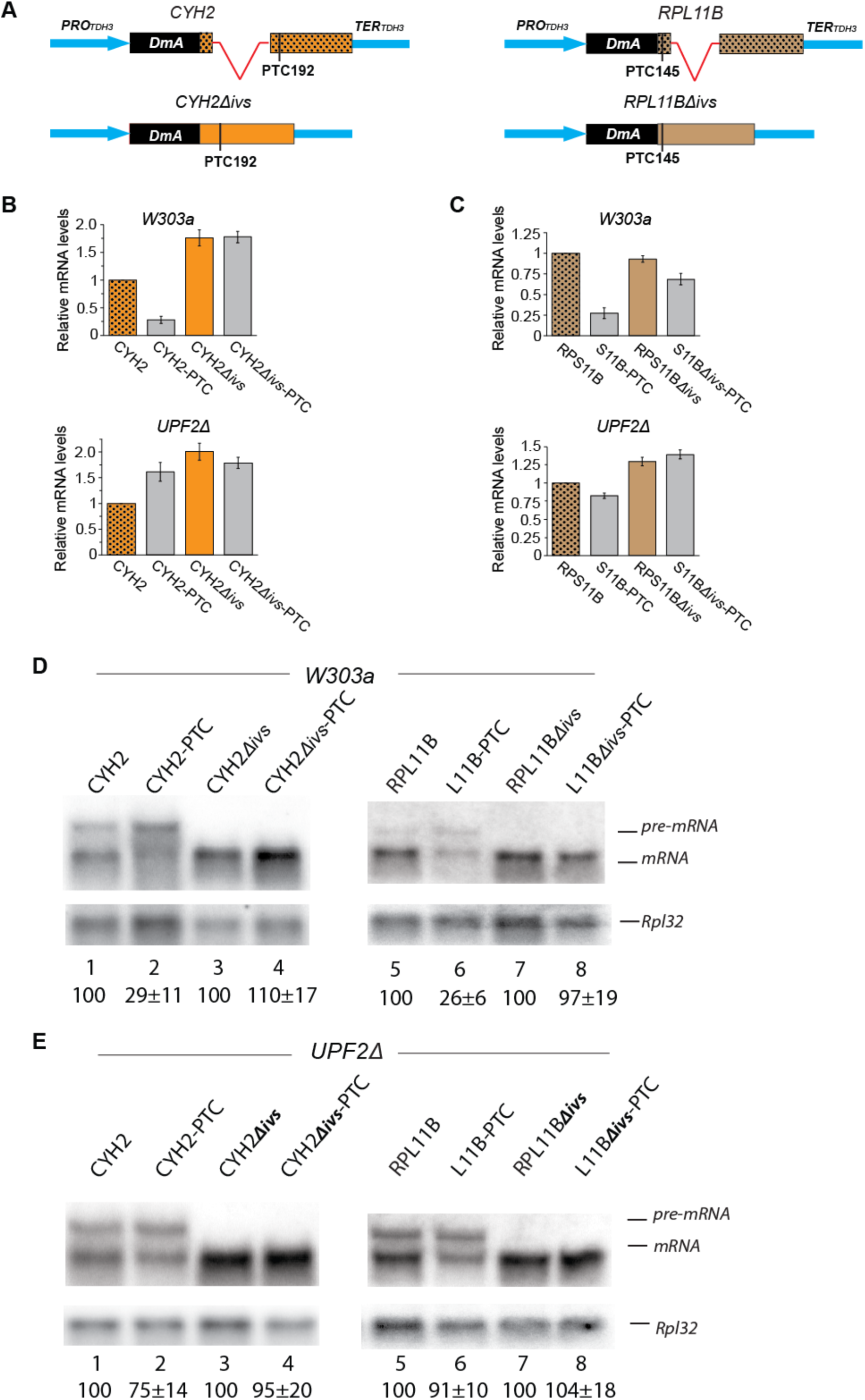
Splicing enhanced NMD in endogenous genes. (**A**) Schematics of NMD reporters expressing two endogenous intron-containing genes, *CYH2* and *RpL11B*, and corresponding cDNA derivatives, lacking the intron. A 300 bp DNA fragment from the *Drosophila melanogaster Adh* gene (*DmA*) was fused to the 5’end of the constructs to allow detection by Northern blotting. (**B**) The steady-state mRNA levels of *CYH2* reporters, quantified by real-time PCR in wild-type (top panel) and *UPF2Δ* (bottom panel). CYH2-PTC refers to the construct with a PTC (TAA) at codon position 192 of the coding region (62% of the sequence downstream from the start codon) which is located 54 nt downstream of the intron. CYH2*Δivs* indicated the reporter where the intron was removed. (**C**) Results of similar experiments, as described in B, but using the *RPS11B* gene-derived reporters, in either *W303a* (top) or *UPF2Δ* (bottom). S11B-PTC refers to the construct containing a PTC (TAA) at codon position 145 in the coding region (49% of the coding region), located 30 nt upstream of the intron. RPS11B*Δivs* indicates the intron-less version of the reporter. Measurements are derived from three independent cultures. (**D**) Northern blotting detection of *CYH2* (top left panel) and RPL11B transcripts (top right panel) in wild-type (W303a); corresponding bottom panels show RpL32 mRNA. **(E)** Similar analysis as D, but in *UPF2Δ*. Quantifications are based on three biological repeats.

### An intron must be relatively close to a PTC for splicing to enhance NMD

We examined whether the effect an intron has on NMD depends on its distance from a PTC. Two additional constructs were tested in which a spacer of either 147 bp or 291 bp was inserted at codon position 120, to lengthen the distance between the intron and the PTC (Figure 5A). We found by both RT-PCR quantification (Figure 5B) and Northern blotting (Figure 5C) that the presence of either spacer suppresses the ability of the intron to enhance NMD. Whilst the mRNA level of the original reporter (PTC140-Aivs) was strongly affected by NMD, that of both spacer-containing constructs is unaffected by the PTC and their levels do not increase in *UPF2Δ*, unlike that of PTC140ivsA (Figure 5B and 5C, bottom panels). Consistent with the absence of NMD, the GFPivsA-147 and PTC140ivsA-147 mRNAs have similar stabilities in wild-type, unlike PTC140-Aivs, which is apparently unstable compared to the PTC-less control (Figure 5D); the four transcripts have similar decay profiles in *UPF2Δ* strain (Figure 5E). Cumulatively, the data indicate that an intron must be relatively close to a PTC for splicing to enhance NMD. The distance between the intron and the PTC is 90 nt in the NMD-sensitive construct, compared to 243 nt and 387 nt in the two NMD-insensitive constructs described just above. Although the mRNA levels of the spacer-containing reporters are lower than those without a spacer, this, as well as the lack of effect of the intron on NMD, is unlikely to be due to either spacer reducing splicing efficiency. The pre-mRNA levels are similar among all constructs (Figure 5C). The longer pre-mRNAs could potentially be more affected by NMD as either spacer lengthens the distance between the intron-encoded PTC and the normal 3’UTR, however, their levels are also similar to those of the constructs without a spacer in *UPF2Δ*, consistent with all three pre-mRNAs being equally affected by NMD (Figure 5E). Our observations are consistent with the previous report that a spacer that lengthens the downstream exon by 250 bp does not change splicing efficiency in *S. cerevisiae* (Tardiff et al., 2006). Moreover, regardless of whether the inserts could affect splicing, or more likely other processes, and results in less mRNA, the conclusion that lengthening the distance of the intron from the PTC suppresses NMD, also in this instance is solely drawn from pair-wise comparisons of mRNA levels of PTC+ vs PTC-less version of otherwise identical reporters. Additionally, the translation yields of both GFPivsA-147 and ivsA-291 are similar, or moderately lower than the initial spacer-less GFP-Aivs reporter when assessed by Western blotting, in both wild-type and *UPF2Δ* (Supplementary Fig 2), suggesting that the suppression effect that either spacer has on NMD is not a consequence of decreased translation efficiency of the transcripts

**Figure 5.**
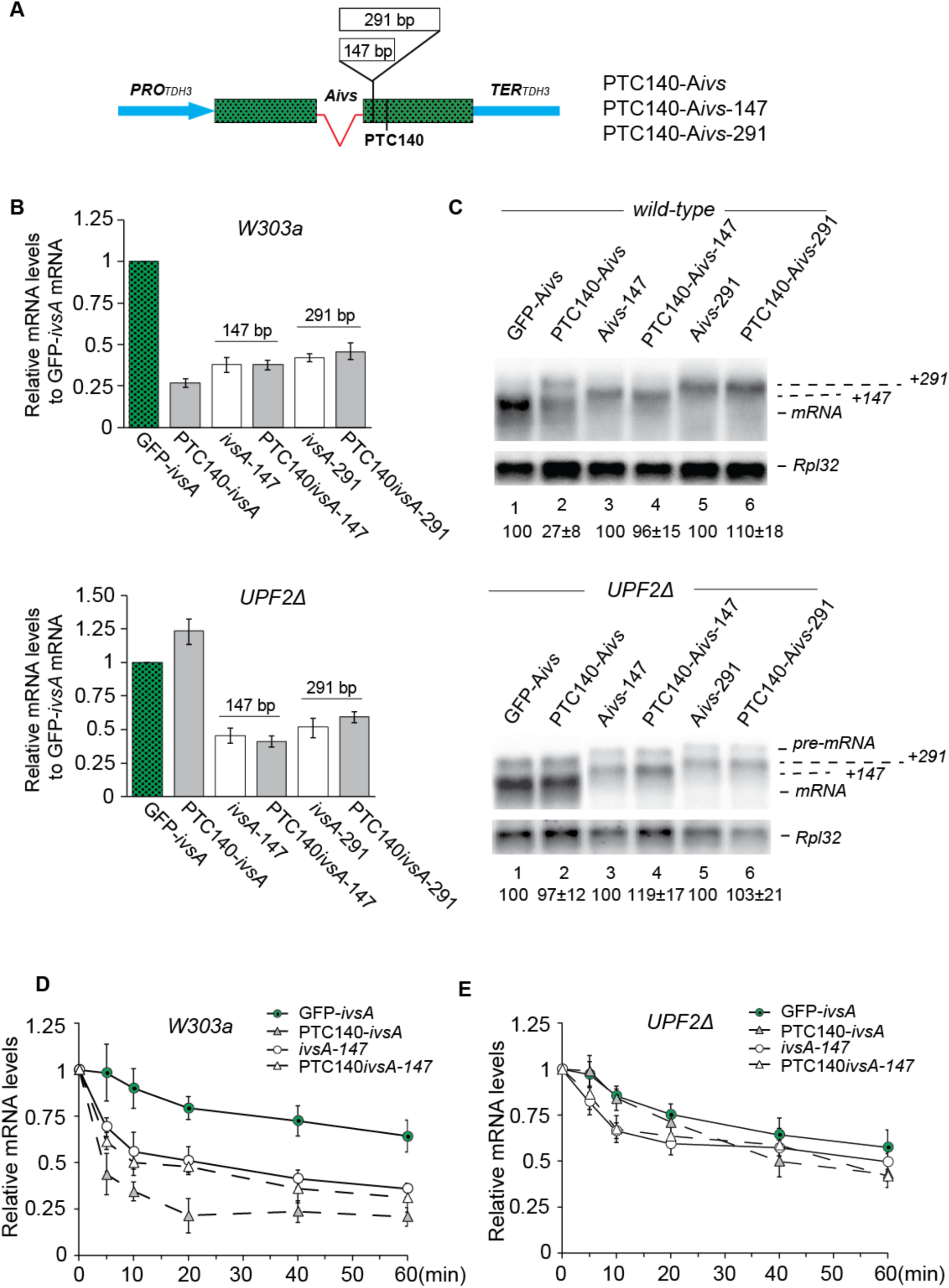
Lengthening the distance between an intron and a PTC abolishes splicing-dependent NMD. (**A**) Schematic maps of NMD reporters carrying different spacer inserts between PTC140 and the intron. The two spacers, 147 bp and 291 bp from luciferase gene, were inserted at codon position 120. (**B**) RT-PCR quantification of steady-state mRNA levels from the reporters relative to GFP-*ivsA*, in *W303a* (B) and *UPF2Δ* (C). (**C**) Northern blotting analysis of GFP transcripts (top panel) in W303a and *UPF2Δ* (bottom); bottom images show the *Rpl32* signal used as a loading control. (**D and E**) Decay profiles of the different mRNAs calculated by quantitative RT-PCR at different time points after transcription block, as detailed in Supplementary Figure 1. All quantifications are based on three biological repeats.

### An intron effect on translation yield does not correlate to its effect on NMD

To examine further whether the NMD-enhancing effect of splicing might be an indirect consequence of a change in translation efficiency of spliced mRNAs, we generated reporters in which the firefly luciferase coding sequence was fused to that of GFP, which was either intron-less or carrying Aivs or Civs (Figure 6A). These were transformed into the wild-type, NMD mutants. To estimate translation yield, luciferase activity and mRNA levels were quantified in parallel (Figure 6B and 6C). The spliced A-*ivs* mRNA appears to be translated more efficiently than the intron-less reporter in wild-type and some of the mutant strains, yet translation of the C-*ivs* mRNA was only slightly increased (Figure 6B), despite the two introns enhancing NMD to similar extents. Therefore, spliced mRNAs might be translated more efficiently, as reported previously in yeast and other organisms (Shaul, 2017); however, our data indicate that this effect does not correlate with either the extent of NMD, or its suppression in the NMD mutant strains.

**Figure 6.**
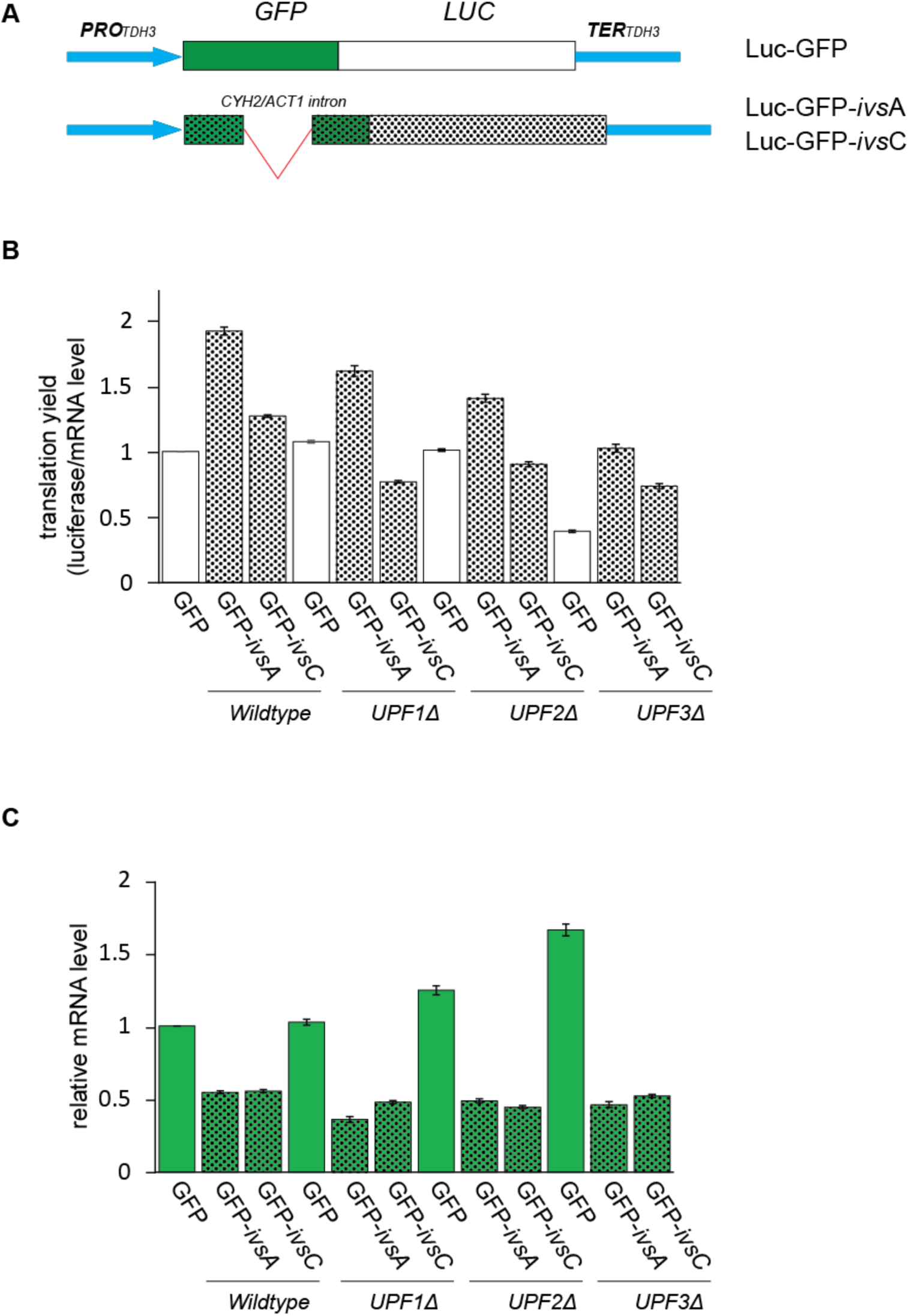
Spliced mRNAs are as efficient translated as non-spliced mRNAs in *PRP17Δ*. (**A**) Diagram of the GFP reporters: intron-less GFP (top), GFP containing either *ACT1* or *CYH2* intron (below), both fused with firefly luciferase at the carboxyl terminal. (**B**). The three plasmid constructs were transformed into either wild type (*BY4742), UPF1Δ, UPF2Δ, UPF3Δ, PRP17Δ, CWC21Δ* or *LSM6Δ*. Relative levels of GFP mRNA were quantified by qRT-PCR using primers that span the exon junction making them specific for the spliced transcript, from equal amounts of total-RNA (detailed in Supplementary Figure 1). Quantifications are based on three separate experiments. (**C**) Luciferase activities measured in aliquots of the same cultures used for the mRNA quantification in B. Luciferase relative light units were normalized to mRNA levels and are shown as means of three measurements with standard deviations.

## Discussion

Contrasting expectations that NMD is independent of splicing in *S. cerevisiae*, this investigation reveals that inserting an intron nearby a PTC also strongly enhances NMD in this organism. We examined constructs containing different introns, either upstream or downstream of a PTC and found that an intron could enhance NMD regardless of whether it was located before or after the PTC. The data also indicate that an intron must be relatively close to a PTC to enhance NMD. Apart from the proximity to the PTC, another determinant could be intron length. Of the three introns examined, the most effective at enhancing NMD were the long *ACT1* and *CYH2* introns, whereas the short *UBC4* intron had only a moderate enhancing effect on NMD even though it was the most efficiently spliced of the three. This effect of splicing on NMD has possibly not been previously recognized in *S. cerevisiae* because there are only 10 genes with more than one intron in this organism, all of which have two introns (Hossain et al., 2011). Notably though, in *SUS1*, which carries two introns, the level of an alternatively spliced transcript, where the first intron is retained and the second spliced, was previously reported to be increased in *UPF1Δ*, whilst this was not the case for the transcript in which both introns are retained (Hossain et al., 2011). In view of the data we have presented, it is plausible that splicing of the second intron triggers NMD of the alternatively spliced *SUS1* transcript, as the retention of its first intron introduces a PTC (see sequence in Saccharomyces Genome Database), which does not seem to induce apparent splicing-independent NMD as reported for unspliced transcript of highly expressed ribosomal protein genes which characteristically have a single intron at the beginning of the coding region (He et al., 1993; Sayani et al., 2008).

We also find that both the *ACT1* and *CYH2* introns enhance translation yield, as previously reported in this and many other eukaryotes. It is therefore plausible that the effect on NMD is a consequence of increased ribosome scanning – introns enhance translation yield in many eukaryotes (Shaul, 2017). However, we find no clear correlation between increased translation yield and the extent to which these introns enhance NMD. Moreover, higher translation efficiency correlates with reduced rather than higher NMD-sensitivity in *S. cerevisiae* (Celik et al., 2017). An EJC-like model, even if molecularly distinct, perhaps involving non-EJC splicing factors as previously reported in other organisms (Aznarez et al., 2018; Michlewski et al., 2008), remains a possible explanation for these observations. However, an EJC-like model still cannot explain how an intron could enhance NMD when located upstream of the PTC. Many such complexes associated with the mRNA upstream of a PTC will have to be removed by the ribosome before reading the PTC, as the tight diameter of the ribosomal mRNA tunnel allows only unstructured RNA deprived of associated proteins to pass through (Qin et al., 2014; Yusupova et al., 2001).

Finally, our observations suggest that lack of adjacent flanking introns might be a reason why in *S. cerevisiae*, unlike in mammalian cells for instance, only PTCs located early in the coding region typically induce strong NMD. This “polarity effect” has been described since the first report of NMD and well-documented in subsequent studies (Losson and Lacroute, 1979; Peltz et al., 1994). The effect is also apparent in other organisms, for example in the *Adh* gene of *Drosophila* where only PTCs located early in the coding region cause strong NMD (Behm-Ansmant et al., 2007; Brogna, 1999). This strong splicing-independent NMD associated with early PTCs might be mechanistically distinct from NMD triggered by intron proximity to late PTCs which otherwise do not reduce mRNA levels. Based on these and our earlier observations in *S. pombe* (Wen and Brogna, 2010), it is feasible that a determining factor of splicing-independent NMD is the proximity of the PTC from the translation start codon.

## Materials and Methods

### Plasmids and strains

GFP-based NMD reporters were generated by cloning the coding sequence of the S65T GFP variant (GenBank: BD062761) into the BamHI sites of either the pRS416 or pRS304 vector, flanked by TDH3 promoter and terminator sequences (Kuperwasser et al., 2004). Mutations were introduced by overlapping PCR and verified by sequencing. The intron of the pGFP-ivs construct is derived from the *ACT1, CYH2* or *UBC4* gene, PCR amplified from *S. cerevisiae* genomic DNA and cloned into the PmlI site at position 328 of the coding region (codon 110, out of frame) by blunt-end ligation. The pGFP-A*ivs*-147 and pGFP-A*ivs*-291 constructs were generated by inserting either a 147 bp or a 291 bp DNA fragment in the AvrII site by overlapping PCR; AvrII site is 34 bp downstream of the *ACT1* intron. Both 147 bp and 291 bp DNA inserts correspond to overlapping regions of the firefly luciferase gene (see primer sequences in Supplementary Table 2). The *RpL11B* and the *CYH2* genes were amplified from either yeast genomic DNA or cDNA while nonsense mutations were introduced by overlapping PCR and then cloned into pRS304-*Dm*ADH (see primer sequences in Supplementary Table 2). The main strains used in the NMD assessment are listed in Supplementary Table 1.

### Yeast transformation

Transformations were performed using the lithium acetate method as previously described, with minor modifications (Soni et al., 1993). In brief, 100 μL of yeast cell pellet (about 10^8^ cells per transformation) was resuspended and vigorously vortexed in 360 μL of freshly prepared transformation mix (33% PEG 3350, 100 μg ssDNA, 500 ng plasmid DNA in 0.1 M LiAc), incubated at 30°C for 20 minutes, and then heat shocked at 42°C in a water bath for 45 minutes. After heat shock, the yeast cells were pelleted by centrifugation, at 5000 rpm for 1 min, washed once with 1 mL of sterile water, resuspended in 1 mL of the same, after which 1/10 to 1/5 was spread on the selective Synthetic Drop-out Media plate (Formedium, UK). The plates were typically incubated at 30°C for 2 days until colonies of the transformants reached up to 0.5 mm in diameter.

### Northern blot assay

Total RNA was typically extracted from 10 mL cell cultures grown to 0.8-1 OD_600_, using the hot acid phenol method, as described (Ausubel et al., 1996). RNA was separated on 1.2% agarose gels in the presence of formaldehyde. RNA was blotted overnight by capillary transfer onto a nylon membrane (Hybond-N, GE Healthcare) and hybridized with ^32^P random priming-labelled probes, as described (Yang et al., 1993). Probes were PCR amplified from plasmid clones (*GFP* and *DmADH*) or from genomic DNA (*rpL32*). Radioactive membranes were imaged using a phosphorimager (Molecular Image™-FX, Bio-Rad), and the intensity of the bands was calculated using the Quantity One software (Bio-Rad).

### RT-PCR RNA levels and stability quantification

First-strand cDNA was synthesized using a PrimeScript™ RT reagent kit (Takara, China) from total-RNA according to manufacturer instructions. RNA was first treated by 1 Unit DNase I (RNase free, TaKaRa, China) at 37.0 °C for 20 min. Real-time PCR was performed using a Bio-Rad CFX96 system (Bio-Rad, Hercules, CA, UA) according to the manufacturer guidance. Reactions were performed in 25 μL containing SYBR Green I dye and 0.5 μL cDNA. 18S rRNA (1/100 dilution of RT product) was used as the internal reference for RNA level normalization. The following cycling parameters were applied, 95.0 °C for 2 min followed by 40 cycles at 95.0 °C for 10 s, 55.0 °C for 15 s, and 72.0 °C for 30 s. The 2^−ΔCT^ method was used to calculate relative expression levels of the target transcripts and normalized to 18S rRNA, and all samples were analyzed in triplicate. The primers used are listed in supplementary Table 2. To block transcription, strains were cultured to OD_600_ 0.5-0.6 and treated with 150 μg/mL 1, 10 phenanthroline at time zero and then transferred in the tube half-filled with ice at different time points, prior RNA extraction using the hot phenol as described above.

### Luciferase assay

Yeast culture densities were adjusted to 1 OD_600_. A 10 μL of each culture was then mixed with 15 μL of sterile purified water and loaded onto a 96-well microtiter plate, which contained 25 μL of Steady-Glo luciferase substrate solution (Promega) in each well. Chemiluminescence was measured in triplicates of each culture using a microplate reader (Tecan). Relative GFP mRNA levels were quantified by real-time qRT-PCR with primers spanning the exon junction specific for the spliced transcript (GFP.Qs.rev and GFP.Q.for, Supplementary Table 2), from equal amounts of total-RNA.

## Acknowledgments

We thank Stephen Dove for help and support, Michael Rosbash, He Feng, Alan Jacobson and Jean D. Beggs for providing reagents. This work was supported by the Wellcome Trust (9340/Z/09/Z) and BBSRC (BB/M022757/1) project grants to SB, and a grant from National Natural Science Foundation of China (NSFC, Grant No. 34171234) to JW.

## Contributions

Saverio Brogna (SB) and Jikai Wen (JW) conceived the study. JW performed most of the experiments. JW and LM performed the mutants screening. MH performed the mRNA decay measurements. JW, MP, AS, VD and Jianming Wang performed the polysomal fractionation and Western blots. SB and JW wrote the paper. KP contributed technical assistance. TM helped write the paper.

## References

Abelson, J., Blanco, M., Ditzler, M.A., Fuller, F., Aravamudhan, P., Wood, M., Villa, T., Ryan, D.E., Pleiss, J.A., Maeder, C., et al. (2010). Conformational dynamics of single pre-mRNA molecules during in vitro splicing. Nat Struct Mol Biol 17, 504–512.

Ares, M., Grate, L., and Pauling, M.H. (1999). A handful of intron-containing genes produces the lion’s share of yeast mRNA. Rna-a Publication of the Rna Society 5, 1138–1139.

Ausubel, F.A., Brent, R., Kingston, R.E., Moore, D.D., Seidman, J.G., Smith, J.A., and K., S. (1996). Current Protocols in Molecular Biology. (New York: John Wiley & Sons).

Aznarez, I., Nomakuchi, T.T., Tetenbaum-Novatt, J., Rahman, M.A., Fregoso, O., Rees, H., and Krainer, A.R. (2018). Mechanism of Nonsense-Mediated mRNA Decay Stimulation by Splicing Factor SRSF1. Cell Reports 23, 2186–2198.

Behm-Ansmant, I., Gatfield, D., Rehwinkel, J., Hilgers, V., and Izaurralde, E. (2007). A conserved role for cytoplasmic poly(A)-binding protein 1 (PABPC1) in nonsense-mediated mRNA decay. Embo J 26, 1591–1601.

Brogna, S. (1999). Nonsense mutations in the alcohol dehydrogenase gene of Drosophila melanogaster correlate with an abnormal 3’ end processing of the corresponding pre-mRNA. RNA 5, 562–573.

Brogna, S., McLeod, T., and Petric, M. (2016). The Meaning of NMD: Translate or Perish. Trends Genet 32, 395–407.

Brogna, S., and Wen, J. (2009). Nonsense-mediated mRNA decay (NMD) mechanisms. Nat Struct Mol Biol 16, 107–113.

Buhler, M., Steiner, S., Mohn, F., Paillusson, A., and Muhlemann, O. (2006). EJC-independent degradation of nonsense immunoglobulin-mu mRNA depends on 3’ UTR length. Nat Struct Mol Biol 13, 462–464.

Carter, M.S., Li, S.L., and Wilkinson, M.F. (1996). A splicing dependent regulatory mechanism that detects translation signals. Embo J 15, 5965–5975.

Celik, A., Baker, R., He, F., and Jacobson, A. (2017). High-resolution profiling of NMD targets in yeast reveals translational fidelity as a basis for substrate selection. RNA 23, 735–748.

Eberle, A.B., Stalder, L., Mathys, H., Orozco, R.Z., and Muhlemann, O. (2008). Posttranscriptional gene regulation by spatial rearrangement of the 3’ untranslated region. PLoS Biol 6, e92.

Fatscher, T., Boehm, V., and Gehring, N.H. (2015). Mechanism, factors, and physiological role of nonsense-mediated mRNA decay. Cell Mol Life Sci 72, 4523–4544.

Gatfield, D., Unterholzner, L., Ciccarelli, F.D., Bork, P., and Izaurralde, E. (2003). Nonsense-mediated mRNA decay in Drosophila: at the intersection of the yeast and mammalian pathways. Embo J 22, 3960–3970.

He, F., and Jacobson, A. (2015). Nonsense-Mediated mRNA Decay: Degradation of Defective Transcripts Is Only Part of the Story. Annu Rev Genet.

He, F., Peltz, S.W., Donahue, J.L., Rosbash, M., and Jacobson, A. (1993). Stabilization and ribosome association of unspliced pre-mRNAs In a yeast Upf1-mutant. Proc Natl Acad Sci U S A 90, 7034–7038.

Hossain, M.A., Rodriguez, C.M., and Johnson, T.L. (2011). Key features of the two-intron Saccharomyces cerevisiae gene SUS1 contribute to its alternative splicing. Nucleic Acids Res 39, 8612–8627.

Hug, N., Longman, D., and Caceres, J.F. (2016). Mechanism and regulation of the nonsense-mediated decay pathway. Nucleic Acids Res 44, 1483–1495.

Hurt, J.A., Robertson, A.D., and Burge, C.B. (2013). Global analyses of UPF1 binding and function reveal expanded scope of nonsense-mediated mRNA decay. Genome Res 23, 1636–1650.

Jaffrey, S.R., and Wilkinson, M.F. (2018). Nonsense-mediated RNA decay in the brain: emerging modulator of neural development and disease. Nat Rev Neurosci 19, 715–728.

Juneau, K., Miranda, M., Hillenmeyer, M.E., Nislow, C., and Davis, R.W. (2006). Introns regulate RNA and protein abundance in yeast. Genetics 174, 511–518.

Karousis, E.D., and Muhlemann, O. (2019). Nonsense-Mediated mRNA Decay Begins Where Translation Ends. Cold Spring Harb Perspect Biol 11.

Kawashima, T., Douglass, S., Gabunilas, J., Pellegrini, M., and Chanfreau, G.F. (2014). Widespread use of non-productive alternative splice sites in Saccharomyces cerevisiae. PLoS genetics 10, e1004249.

Kawashima, T., Pellegrini, M., and Chanfreau, G.F. (2009). Nonsense-mediated mRNA decay mutes the splicing defects of spliceosome component mutations. Rna-a Publication of the Rna Society 15, 2236–2247.

Kuperwasser, N., Brogna, S., Dower, K., and Rosbash, M. (2004). Nonsense-mediated decay does not occur within the yeast nucleus. RNA 10, 1907–1915.

Kupfer, D.M., Drabenstot, S.D., Buchanan, K.L., Lai, H.S., Zhu, H., Dyer, D.W., Roe, B.A., and Murphy, J.W. (2004). Introns and splicing elements of five diverse fungi. Eukaryot Cell 3, 1088–1100.

Kurosaki, T., Popp, M.W., and Maquat, L.E. (2019). Quality and quantity control of gene expression by nonsense-mediated mRNA decay. Nat Rev Mol Cell Biol 20, 406–420.

Le Hir, H., Izaurralde, E., Maquat, L.E., and Moore, M.J. (2000a). The spliceosome deposits multiple proteins 20-24 nucleotides upstream of mRNA exon-exon junctions. Embo J 19, 6860–6869.

Le Hir, H., Moore, M.J., and Maquat, L.E. (2000b). Pre-mRNA splicing alters mRNP composition: evidence for stable association of proteins at exon-exon junctions. Genes Dev 14, 1098–1108.

Le Hir, H., Sauliere, J., and Wang, Z. (2016). The exon junction complex as a node of post-transcriptional networks. Nat Rev Mol Cell Biol 17, 41–54.

Leeds, P., Wood, J.M., Lee, B.S., and Culbertson, M.R. (1992). Gene products that promote mRNA turnover in Saccharomyces cerevisiae. Mol Cell Biol 12, 2165–2177.

Lesser, C.F., and Guthrie, C. (1993). Mutational analysis of pre-mRNA splicing in Saccharomyces cerevisiae using a sensitive new reporter gene, CUP1. Genetics 133, 851–863.

Lin, R.J., Newman, A.J., Cheng, S.C., and Abelson, J. (1985). Yeast mRNA splicing in vitro. J Biol Chem 260, 14780–14792.

Lindeboom, R.G.H., Supek, F., and Lehner, B. (2016). The rules and impact of nonsense-mediated mRNA decay in human cancers. Nature Genetics 48, 1112–1118.

Lloyd, J.P.B. (2018). The evolution and diversity of the nonsense-mediated mRNA decay pathway. F1000Res 7, 1299.

Losson, R., and Lacroute, F. (1979). Interference of nonsense mutations with eukaryotic messanger RNA stability. Proc Natl Acad Sci USA 76, 5134–5137.

Lykke-Andersen, S., and Jensen, T.H. (2015). Nonsense-mediated mRNA decay: an intricate machinery that shapes transcriptomes. Nat Rev Mol Cell Biol 16, 665–677.

Maquat, L.E. (2004). Nonsense-mediated mRNA decay: splicing, translation and mRNP dynamics. Nat Rev Mol Cell Biol 5, 89–99.

Mendell, J.T., Sharifi, N.A., Meyers, J.L., Martinez-Murillo, F., and Dietz, H.C. (2004). Nonsense surveillance regulates expression of diverse classes of mammalian transcripts and mutes genomic noise. Nat Genet 36, 1073–1078.

Metzstein, M.M., and Krasnow, M.A. (2006). Functions of the nonsense-mediated mRNA decay pathway in Drosophila development. PLoS genetics 2, e180.

Michlewski, G., Sanford, J.R., and Caceres, J.F. (2008). The splicing factor SF2/ASF regulates translation initiation by enhancing phosphorylation of 4E-BP1. Mol Cell 30, 179–189.

Nagy, E., and Maquat, L.E. (1998). A rule for termination-codon position within intron-containing genes: when nonsense affects RNA abundance. Trends Biochem Sci 23, 198–199.

Nasif, S., Contu, L., and Muhlemann, O. (2018). Beyond quality control: The role of nonsense-mediated mRNA decay (NMD) in regulating gene expression. Semin Cell Dev Biol 75, 78–87.

Nyiko, T., Kerenyi, F., Szabadkai, L., Benkovics, A.H., Major, P., Sonkoly, B., Merai, Z., Barta, E., Niemiec, E., Kufel, J., et al. (2013). Plant nonsense-mediated mRNA decay is controlled by different autoregulatory circuits and can be induced by an EJC-like complex. Nucleic Acids Res 41, 6715–6728.

Peltz, S.W., He, F., Welch, E., and Jacobson, A. (1994). Nonsense-mediated mRNA decay in yeast. Prog Nucleic Acid Res Mol Biol 47, 271–298.

Qin, P.W., Yu, D.M., Zuo, X.B., and Cornish, P.V. (2014). Structured mRNA induces the ribosome into a hyper-rotated state. EMBO reports 15, 185–190.

Sayani, S., Janis, M., Lee, C.Y., Toesca, I., and Chanfreau, G.F. (2008). Widespread impact of nonsense-mediated mRNA decay on the yeast intronome. Mol Cell 31, 360–370.

Schreiber, K., Csaba, G., Haslbeck, M., and Zimmer, R. (2015). Alternative Splicing in Next Generation Sequencing Data of Saccharomyces cerevisiae. Plos One 10.

Shaul, O. (2017). How introns enhance gene expression. Int J Biochem Cell B 91, 145–155.

Silva, A.L., Ribeiro, P., Inacio, A., Liebhaber, S.A., and Romao, L. (2008). Proximity of the poly(A)-binding protein to a premature termination codon inhibits mammalian nonsense-mediated mRNA decay. RNA 14, 563–576.

Singh, A.K., Choudhury, S.R., De, S., Zhang, J., Kissane, S., Dwivedi, V., Ramanathan, P., Orsini, L., Hebenstreit, D., and Brogna, S. (2019). The RNA helicase UPF1 associates with mRNAs cotranscriptionally and is required for the release of mRNAs from transcription sites. Elife.

Singh, G., Rebbapragada, I., and Lykke-Andersen, J. (2008). A competition between stimulators and antagonists of Upf complex recruitment governs human nonsense-mediated mRNA decay. PLoS Biol 6, e111.

Soni, R., Carmichael, J.P., and Murray, J.A. (1993). Parameters affecting lithium acetate-mediated transformation of Saccharomyces cerevisiae and development of a rapid and simplified procedure. Curr Genet 24, 455–459.

Swida, U., Thuroff, E., and Kaufer, N.F. (1986). Intron Mutations That Affect the Splicing Efficiency of the *CYH2* Gene of Saccharomyces-Cerevisiae. Molecular & General Genetics 203, 300–304.

Tardiff, D.F., Lacadie, S.A., and Rosbash, M. (2006). A genome-wide analysis indicates that yeast pre-mRNA splicing is predominantly posttranscriptional. Mol Cell 24, 917–929.

Thermann, R., NeuYilik, G., Deters, A., Frede, U., Wehr, K., Hagemeier, C., Hentze, M.W., and Kulozik, A.E. (1998). Binary specification of nonsense codons by splicing and cytoplasmic translation. Embo J 17, 3484–3494.

Tian Miao, W.Y., Jing Zhang, Huai Dang, Xingyi Lu, Chengjie Fu, Miao Wei (2017). Nonsense-mediated mRNA decay in Tetrahymena is EJC independent and requires a protozoa-specific nuclease. Nucleic Acids Res gkx256.

Trcek, T., Sato, H., Singer, R.H., and Maquat, L.E. (2013). Temporal and spatial characterization of nonsense-mediated mRNA decay. Genes Dev 27, 541–551.

Wang, J., Gudikote, J.P., Olivas, O.R., and Wilkinson, M.F. (2002). Boundary-independent polar nonsense-mediated decay. EMBO reports 3, 274–279.

Wen, J., and Brogna, S. (2010). Splicing-dependent NMD does not require the EJC in Schizosaccharomyces pombe. The Embo Journal 29, 1537–1551.

Yang, H., McLeese, J., Weisbart, M., Dionne, J.L., Lemaire, I., and Aubin, R.A. (1993). Simplified high throughput protocol for northern hybridization. Nucleic Acids Res 21, 3337–3338.

Yusupova, G.Z., Yusupov, M.M., Cate, J.H., and Noller, H.F. (2001). The path of messenger RNA through the ribosome. Cell 106, 233–241.

Zhang, J., and Maquat, L.E. (1996). Evidence That the Decay Of Nucleus-Associated Nonsense Messenger-Rna For Human Triosephosphate Isomerase Involves Nonsense Codon Recognition After Splicing. Rna-a Publication Of the Rna Society 2, 235–243.

Zhang, J., Sun, X.L., Qian, Y.M., LaDuca, J.P., and Maquat, L.E. (1998). At least one intron is required for the nonsense-mediated decay of triosephosphate isomerase mRNA: a possible link between nuclear splicing and cytoplasmic translation. Mol Cell Biol 18, 5272–5283.

Zhang, Y., and Sachs, M.S. (2015). Control of mRNA Stability in Fungi by NMD, EJC and CBC Factors Through 3’UTR Introns. Genetics 200, 1133-1148.

