## Supplementary tables and figures for "An intron proximal to a PTC enhances NMD in *Saccharomyces cerevisiae*"

(This consists of two tables followed by two figures with legends)

**Supplementary Table 1. Strains used in this study**

| Strain names | genotype | Origin |
|---|---|---|
| W303a | *MATa ade2-1 trp1-1 can1-100 leu2-3,112 his3-11,15 ura3-1* | Gift from Michael Rosbash lab (Dower K., et al., 2004) |
| UPF1Δ | *MATa ade2-1 trp1-1 can1-100 leu2-3,112 his3-11,15 ura3-1 upf1::HIS3* | Gift from Allan Jacobson lab (He F., et al, 2001) |
| UPF2Δ | *MATa ade2-1 trp1-1 can1-100 leu2-3,112 his3-11,15 ura3-1 upf2::HIS3* | Gift from Allan Jacobson lab (He F., et al, 2001) |
| UPF3Δ | *MATa ade2-1 trp1-1 can1-100 leu2-3,112 his3-11,15 ura3-1 upf2::HIS3* | Gift from Allan Jacobson lab (He F., et al, 2001) |
| BY4742 | *MATα his3Δ1 leu2Δ0 lys2Δ0 ura3Δ0* | Gift from Steve Dove's lab |
| UPF1Δ | *MATα his3Δ1 leu2Δ0 lys2Δ0 ura3Δ0 upf1::KanMX6* | Yeast Knock-out (YKO) deletion collection |
| UPF2Δ | *MATα his3Δ1 leu2Δ0 lys2Δ0 ura3Δ0 upf2::KanMX6* | Yeast Knock-out (YKO) deletion collection |
| UPF3Δ | *MATα his3Δ1 leu2Δ0 lys2Δ0 ura3Δ0 upf3::KanMX6* | Yeast Knock-out (YKO) deletion collection |

**Supplementary Table 2. Primers used in this study**

| Primer names | DNA sequence (5' to 3') |
|---|---|
| Act1.ivs.for | GTATGTTCTAGCGCTTGCAC |
| Act1.ivs.rev | CTAAACATATAATATAGCAAC |
| Cyh2.ivs.for | GTATGTAGTTCCATTTGGAAG |
| Cyh2.ivs.rev | CTGTACAAAAAAAATATTGTAATG |
| Uivs.full.for | GTATGTCTAAAGTTATGGCCACGTTTCAAATGCGTGCTTTTTTTTTA AAACTTATGCTCTTATTTACTAACAAAATCAACATGCTATTGAACTA G |
| Uivs.full.rev | CTAGTTCAATAGCATGTTGATTTTGTTAGTAAATAAGAGCATAAGT TTTAAAAAAAAAGCACGCATTTGAAACGTGGCCATAACTTTAGACA TAC |
| ScL11b.P145.for | CCCCGGATCCTCCACTGAA**TAA**ACTGTTCAATCTG |
| ScL11b.wt.for | CCCCGGATCCTCCACTGAATTAACTGTTCAATCTG |
| ScL11b.rev | CCCCGTCGACTTAAAATTTAGCAAATTGCTTGTTGG |
| Sccyh2.wt.for | CCCCGGATCCCCTTCCAGATTCACTAAGACTAGAAAGCAC |
| Sccyh2.rev | CCCCGTCGACTTAAGCGATCAATTCAACAACACC |
| Sccyh2.P192.for | GGTAGAGGTATG**TAA**GGTGGTCAACATCACCACAG |
| Sccyh2.P192.rev | GTTGACCACC**TTA**CATACCTCTACCACCG |
| Sc.l32.for | TACCTCACCCAAAGATTGTC |
| Sc.l32.rev | GGTTTCCAAATCCTTAACG |
| Adh.Bgl.for | AGTCCAGATCTATGTCGTTTACTTTGACCAACAAG |
| Adh.Bam.rev | CTTTGAAGATAAAGAGGATCCAATGTTGCAGATGAT |
| F.Luc.for | CTTGAGCCTAGGTACCCATACGATGTTCCTGA |
| F.Luc147.rev | TTTGTACCTAGGCTCGACCAGGATGGGCAC |
| F.Luc291.rev | TTGGTTCCTAGGGGTCAGGGTGGTCACGAG |
| GFP.Q.for | GAGTTGTCCCAATTCTTGTT |
| GFP.Qs.rev | TTGACTTCAGCACGTGTCTT |
| Aivs.Q.rev | ACAGTTAAATGGGATGGTGC |
| Civs.Q.rev | GTTCTTTCATTCCCTCTTCC |
| 18SrRNA.Q.for | AAACGGCTACCACATCCA |
| 18SrRNA.Q.rev | GCCCAAAGTTCAACTACG |
| RPL32.Q.for | GATTGTCAAGAAGCACACC |
| RPL32.Q.rev | ACCGTAACCGATCTTTGG |
| CYH2.Q.for | CAGAGGTCACGTCTCAGCCG |
| CYH2.Q.rev | GTATTGGTCTCTCTTGTCTTCTGGG |
| S11B.Q.for | AGAGCTTTCCAAAAGCAACCT |
| S11B.Q.rev | CTTGTGTCTCTTTTCGTATCTGTTG |
| GFPas.for | CAACACTTGTCACTACTTTCACTTA |
| GFPas.rev | TTGATTCCATTCTTTTGTTTGTC |

### Supplementary figures and legends

Wen et. al, supplementary figure 1

A

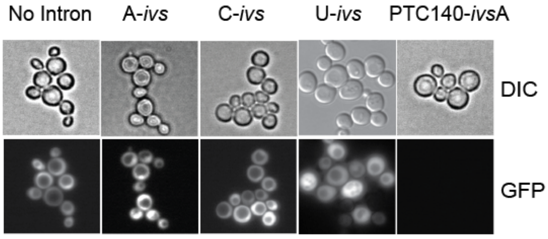


B

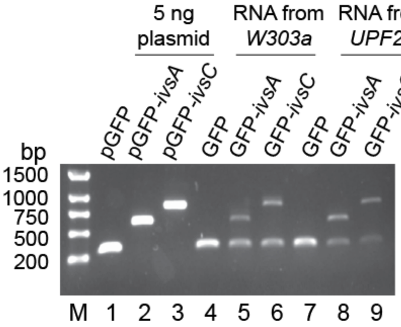


C

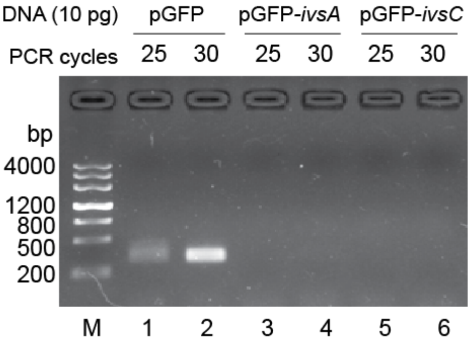


D

Quantitative PCR analysis of DNA by splicing specific primers

| plasmid (10pg) | Cycles |
|---|---|
| pGFP | 23.36 ± 0.24 |
| pGFP-ivsA | N.A. |
| pGFP-ivsC | N.A. |

N.A. Not applicable

E

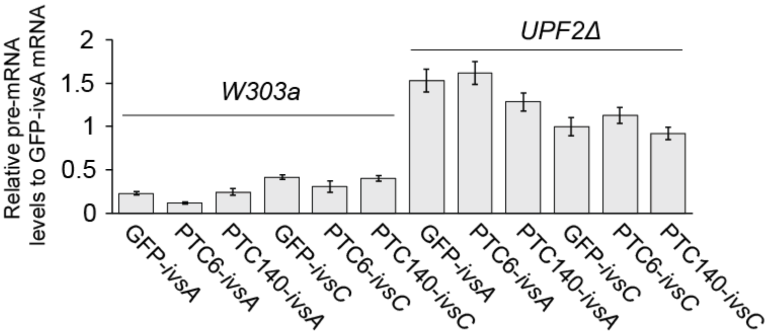


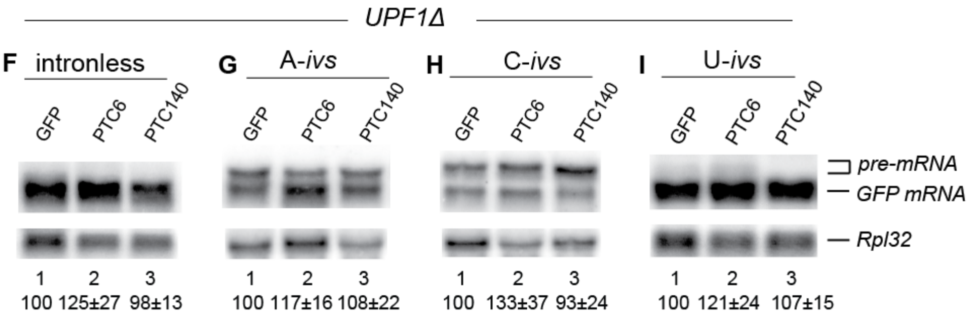

**Supplementary Figure 1. Further validation of the GFP gene reporters.**

**(A)** From left to right – micrographs showing GFP fluorescence of cells transformed with the indicated reporters; the far right panel shows a PTC+ control with no GFP fluorescence. Images were taken with an inverted fluorescence microscope (Nikon Ti). (**B**) PCR and RT-PCR validation of the reporters. Lanes 1-3, PCR amplification of a ~300 bb GFP fragment using primers GFPas.for and GFPas.rev (located ~150 nt at either side of the intron insertion side codon 110) using the indicated plasmids as templates, pGFP (lane 1), pGFP-*ivsA* (lane 2) and pGFP-*ivsC* (lane 3); lanes 4-9, RT-PCR of the GFP transcript with the same GFP primer pair, using total-RNA from either wild-type (lanes 4-6, *W303a*) or *UPF2Δ* (lanes, 7-9) strain transformed with the indicated reporters. The PCRs were run for 25 cycles. (**C** and **D**). Panel C, PCR (25 or 30 cycles) of either the intron-less pGFP plasmid, pGFP-ivsA or pGFP-ivsC with the splicing-specific primer pair (GFP.Q.for and GFP.Qs.rev) used for RT-PCR quantification of the spliced mRNA levels, which only produces the expected fragment with pGFP. (**D**) Ct values of a real-time PCR performed as in C. (**E**) Real-time PCR quantifications of pre-mRNA levels of the indicated NMD reporters, in either wild-type or *UPF2Δ*. Values are expressed as a fraction of the normalized GFP-ivsA mRNA level in wild-type. Both mRNA and pre-mRNA transcript levels were normalized by the 18S rRNA (assessed in parallel using a 1/100 dilution of the same cDNA sample). Pre-mRNAs were detected using a primer in the first exon (GFP.Q.for) and the other in the intron (Aivs.Q.rev or Civs.Q.rev). Values are based on three experimental repeats where error bars indicate standard deviations. (**F-I**) Northern blotting of total-RNA from *UPF1Δ* transformed with the reporters, as indicated. Top panels show hybridization with a GFP-specific probe, the bottom panels show it with a probe specific for the ribosomal protein L32 mRNA (Rpl32), as a loading control. The values below each lane are percentages, with standard deviations, of the level of GFP mRNA relative to that of the corresponding PTC-control, lane 1 in each panel. Quantifications are based on three biological repeats.

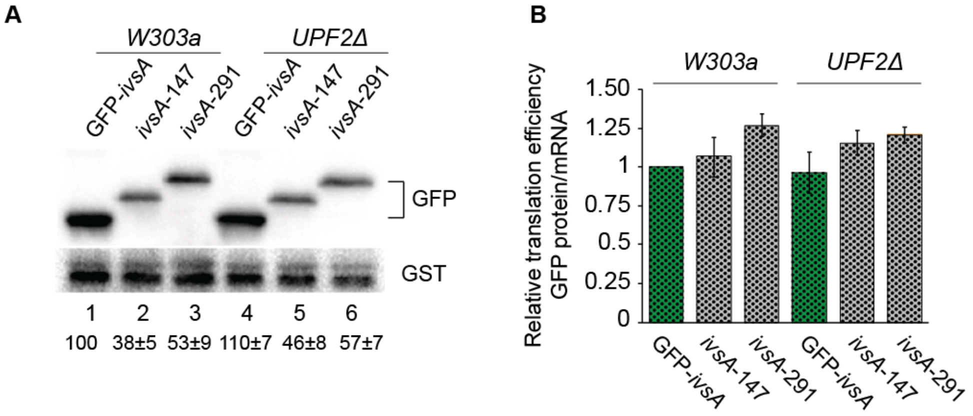


**Supplementary Figure 2. Lengthening of the reporter mRNAs does not reduce their translation yields.**

(**A**) Western blotting of either *W303a* or *UPF2Δ* strain transformed with the indicated GFP expressing reporters. Top panel shows GFP detected with an anti-GFP rabbit polyclonal antibody (Sangon Biotech); bottom panel shows Glutathione S Transferase (GST) protein detected by an anti-GST mouse monoclonal antibody (Sino Biological, 11213-MM01) as the loading control. (**B**) Relative translation yields for the three intron-containing GFP controls, calculated by dividing GFP protein levels by mRNA levels, and expressed as percentages with standard deviations of that produced by GFP-ivs*A* in *W303a* (lane 1). Quantifications are based on three repeats.
